# Would you agree if N is three? On statistical inference for small N

**DOI:** 10.1101/2024.08.26.609821

**Authors:** Eleni Psarou, Christini Katsanevaki, Eric Maris, Pascal Fries

## Abstract

Non-human primate studies traditionally use two or three animals. We previously used standard statistics to argue for using either one animal, for an inference about that sample, or five or more animals, for a useful inference about the population. A recently proposed framework argued for testing three animals and accepting the outcome found in the majority as the outcome that is most representative for the population. The proposal tests this framework under various assumptions about the true probability of the representative outcome in the population, i.e. its typicality. On this basis, it argues that the framework is valid across a wide range of typicalities. Here, we show (1) that the error rate of the framework depends strongly on the typicality of the representative outcome, (2) that an acceptable error rate requires this typicality to be very high (87% for a single type of outlier), which actually renders empirical testing beyond a single animal obsolete, (3) that moving from one to three animals decreases error rates mainly for typicality values of 70-90%, and much less for both lower and higher values. Furthermore, we use conjunction analysis to demonstrate that two out of three animals with a given outcome only allow to infer a lower bound to typicality of 9%, which is of limited value. Thus, the use of two or three animals does not allow a useful inference about the population, and if this option is nevertheless chosen, the inferred lower bound of typicality should be reported.

## Introduction

We have recently argued that a sample of two or few (less than five) subjects allows useful inferences only about that sample but not about the population (Fries & Maris, 2022). On this basis, we recommended to either use a sample of five or more animals to make a useful inference about the population, or to use a sample of one and make an inference about that sample. We discouraged using the traditional approach in non-human primate (NHP) research of using two or three animals, because this leaves the inference limited to the sample while doubling or tripling the number of animals. Note that we are not arguing for any particular number of animals, and certainly not for lowering the statistical standards in the field. To the contrary, we intend to clarify that the traditional use of two or few animals in fixed-effect tests only allows inferences on the sample and thereby much weaker inferences than commonly assumed. If one assumes stronger inferences than actually established with few animals, this can contribute to the replication crisis. This is our first and foremost point. Once this is clarified, we show that even a conjunction test on results from two or few animals, which is typically not performed, would provide only limited gains in inference. Only after these insights, we present our view that these limited gains in inference do not justify the use of a second or third animal, and therefore we recommend to use a single animal. This recommendation is not the starting point, but the conclusion, given the statistical, practical and ethical limits of NHP research.

Soon after we had presented our arguments (Fries & Maris, 2022), a framework was suggested that attempts to draw an inference about the representativeness of an outcome in the population by studying a small number of animals (Laurens, 2022). This framework (1) suggests to “assume that, in each animal, an experiment can lead to a number of qualitatively distinct outcomes”, (2) suggests to “assume a prior distribution across all possible outcomes”, calling “the most likely outcome the ‘representative’ outcome and other ‘outliers’”, (3) considers outlier proportions in the range of 10 to 20%, (4) considers an outcome as representative when the outcome is present in the majority of the tested animals, (5) concludes that requiring two out of three subjects to show an effect strikes an efficient balance between the proportion of correct conclusions and inconclusive outcomes, (6) claims that this conclusion holds “across a wide range of prior distributions” (Laurens, 2022). Here, we critically discuss the proposed framework and conclude that it has serious shortcomings. We note that the term “prior distribution” should not be confused with a prior distribution in a Bayesian framework, and that the framework of Laurens actually does not allow to specify a prior distribution in the Bayesian sense. Most importantly, we show that the framework will only produce an acceptable inference for a narrow range of outcome distributions. We also present a way to estimate a lower bound to the representativeness of an outcome, known as typicality, for a range of M animals tested and N animals showing the outcome. We recommend that this lower bound of typicality is reported in studies that try to draw an inference about the population based on a small number of animals.

## Discussion

### The concept of typicality (also known as prevalence)

The probability of a given test outcome in a population is referred to as the typicality of that outcome in that population (an alternative term is prevalence, which is discussed below in the section on *Additional considerations*). The concept of typicality is central to a technique that has been called conjunction analysis (Friston et al., 1999). A conjunction analysis first tests for a given outcome (e.g. the presence of an effect or a trait) in each subject of a limited sample drawn from a population. It then uses the proportion of the sample with a given outcome to draw an inference about the proportion of the population that would give the same outcome. This proportion of the population is called typicality *γ*. The true value of *γ* cannot be known, but a useful lower bound to typicality, *γ*_*c*_, can be estimated if the false-positive and false-negative rates of the employed tests are specified (Friston et al., 1999), and this is explained in more detail below.

### The N-oo-M framework

The framework proposed by Laurens (2022) starts by testing each investigated subject for a given outcome, and counting in a sample of M subjects the number N of subjects showing that outcome. The framework refers to this as N-out-of-M, or N-oo-M, and refers to e.g. 2 out of 3 tested subjects showing a given outcome as 2-oo-3. After counting the number of animals with a given outcome, the framework aims at drawing a binary inference about which outcome is representative of the population. The framework defines an outcome as representative if it is present in the majority of the tested subjects, i.e. at least two subjects should show the effect in the 2-oo-3 case.

Importantly, the N-oo-M framework suggests to “assume a prior distribution across all possible outcomes”, calling “the most likely outcome the ‘representative’ outcome and other ‘outliers’”. Note that assuming “a prior distribution across all possible outcomes” is not an assumption in the usual sense, because an outcome always has some probability/typicality, and this just follows from the fact that it is a random variable. Conjunction analysis considers one such probability as the parameter of interest and derives a lower bound for this probability/typicality.

The use of the term “prior distribution” may cause confusion among readers that are familiar with Bayesian inference. The framework of Laurens (2022) is not Bayesian, because if it were Bayesian, one would have to specify a prior probability distribution for the parameters, and here this would be typicality *γ* (which in turn specifies the distribution of the outcomes, with probability *γ* for the representative outcome, and 1 − *γ* for the outlier). Because this is a probability, the prior distribution would have a support over the interval [0,1], and usually this is the beta distribution. Combining this prior with the information in the data produces a posterior distribution (via Bayes’ rule) that is a reweighting over the interval [0,1]: segments that were a priori likely/unlikely to contain the true typicality value can be down/upweighted according to the information in the data. Crucially, if this prior were not a distribution over the interval [0,1] but a fixed value (e.g., *γ* = 0.8), there would be no space for down/upweighting. If *γ* is a fixed value and it is known, then no data are required for estimating it. If, on the other hand, *γ* is a fixed unknown value, as in frequentist inference, then data can be used to estimate its value (e.g., by means of a confidence interval). The framework of Laurens (2022) is neither Bayesian nor frequentist; its goal is to estimate the representative outcome, i.e., the event that corresponds to *γ >* 0.5.

### The probability of incorrect conclusions depends on the assumed probability of outliers

If the N-oo-M framework correctly identifies the representative outcome, it defines this as “correct conclusion”. Here, we refer to the probability of correct conclusions as *π*, to the probability of incorrect conclusions (excluding cases considered “inconclusive” by Laurens (2022)) as *δ*, and the probability of outliers as *ω*. The N-oo-M framework suggests specifying the probability of outliers as a “prior” (in the sense of Laurens (2022), see above) and claims that “the N-out-of-M model leads to a similar conclusion across a wide range of prior distributions.” However, Laurens (2022) considers *ω*-values only in a relatively narrow range of 10% to 20%, and allows for percentages to accumulate over different outlier types, including experimental errors. We argue that both, the distributions of outlier probabilities over outlier types, and the total outlier probability, considered by Laurens (2022) are arbitrary.

Fig. 1C of Laurens (2022) shows that with a single type of outlier and with *ω* = 10%, 2-oo-3 reaches *δ* = 2.8%. However, our Fig. 1A shows that for larger values of *ω*, the *δ* increases steeply (light blue curve in our Fig. 1A). Already for an *ω* value of 20%, the *δ* rises to 10.4%, which is more than twice the generally accepted error rate of 5%. Laurens (2022) actually rejected 1-oo-1 at *ω* = 10%, because of *δ* = 10%.

**Figure 1.**
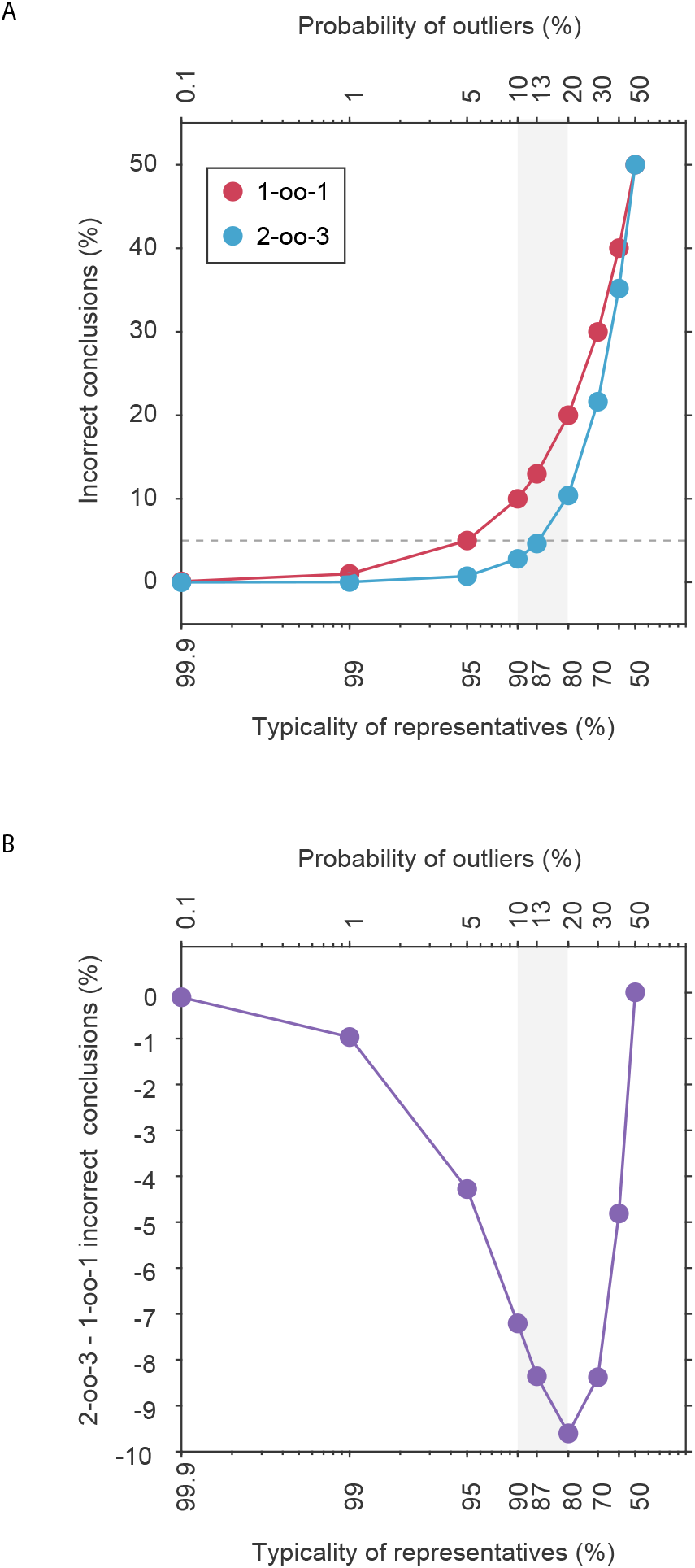
(A) The percentage of incorrect conclusions (*δ*) as a function of the percentage of the assumed typicality of representatives (bottom x-axis) and the assumed probability of outliers (*ω*, top x-axis), for a single type of outliers, and separately for 2-oo-3 (blue) and 1-oo-1 (red). (B) The difference in *δ* for 2-oo-3 versus 1-oo-1. The grey shading indicates the range of outlier proportions considered in Laurens (2022).

Laurens (2022) does actually consider an example with a total *ω* = 20%, yet this *ω* is distributed over three types of outliers occurring at 10%, 5% and 5%, respectively (Fig. 2A of Laurens (2022)), and this specific distribution leads to a *δ* just below 5%, namely at 4.3%. However, we note that this scenario additionally entails a 6.1% possibility of inconclusive cases. Even more worrisome is the fact that this still favorable outcome depends on the precise distribution of *ω* over several outlier types. If a total *ω* of 20% would result from only two types of outliers of 15% and 5%, then *δ* would be 6.8% and thereby higher than the accepted error rate of 5%.

**Figure 2.**
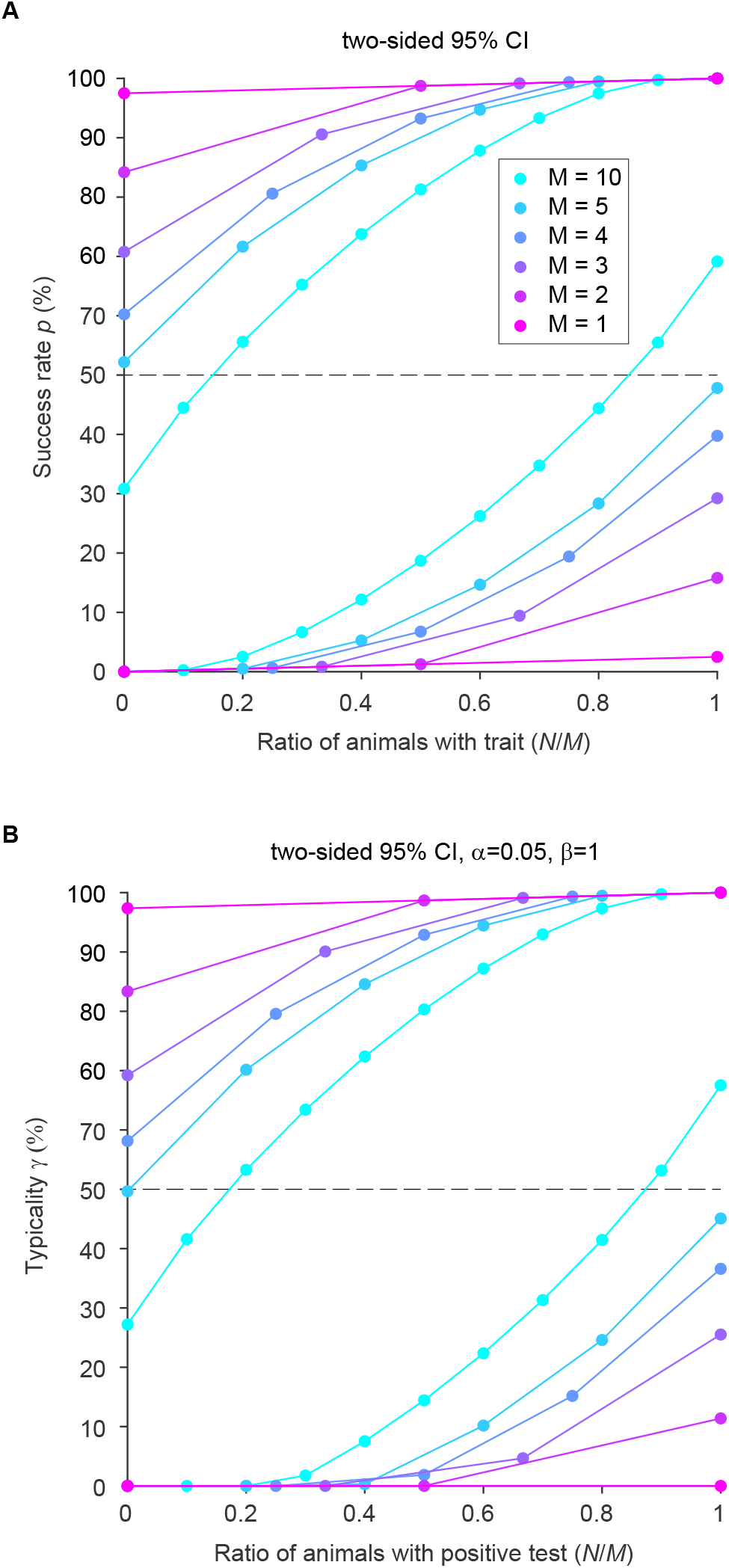
(A) Two-sided 95% confidence intervals (CI) for the success rate (*p*, expressed in percentage) as a function of the ratio of animals with the trait, following Clopper and Pearson (1934). The color legend specifies the different numbers *M* of tested animals. For each *M*, two lines are plotted, corresponding to the upper and the lower limit of the two-sided 95% confidence interval. (B) Same as (A), but for the typicality *γ*.

### A high and immutable prior on the typicality of representatives renders experiments obsolete

Considering all possible combinations of several outlier types is intractable, so we focus in the following on the simple case of one outlier type, yet the reasoning holds for more than one outlier type with correspondingly adjusted numerical results. The blue line in Fig. 1A shows that *δ* starts exceeding the generally accepted error rate of 5% for a probability of outliers (*ω*-value) above 13%, corresponding to a typicality of representatives below 87%. Thus, the 2-oo-3 framework only produces an acceptable error rate for a typicality of representatives of 87% or higher. Here, we point out that if the typicality of representatives has to be >87%, this makes it meaningless to collect empirical data beyond a single animal (one single animal being needed to find out which outcome is actually typical). Remember that the framework aims at a binary decision about which outcome is representative for the population. Yet, for a typicality of representatives >87%, this decision is obsolete.

### 2-oo-3 versus 1-oo-1

The core of the argument presented in favor of the proposed 2-oo-3 framework is a reduction of *δ* for 2-oo-3 compared to 1-oo-1. The *δ* values for 1-oo-1 are shown in Fig. 1A as red line, for 2-oo-3 as blue line, and their difference is shown in Fig. 1B. Here, we point out that the reductions of *δ* that are obtained by moving from 1-oo-1 to 2-oo-3 are a function of *ω*. They peak for *ω* values around 10-30%, close to the ones chosen by Laurens (2022), but they strongly diminish for both larger and smaller *ω* values.

### The problem of defining outliers in small samples

On a general note, it is important to mention that the concept of outliers is problematic in small samples. When a sample is very large, e.g., with 10000 human subjects in a GWAS (genome wide association study), data-driven techniques for outlier detection can be used (Chandola et al., 2009). Such techniques cannot be used if N is two or few. For those very small samples, approaches based on outliers depend on assumptions about the existence and/or likelihood of outliers that are untested.

### Two to three animals allow only limited inferences about the population

The N-oo-M framework can be evaluated through the perspective of conjunction analysis. Conjunction analysis makes no assumption about the typicality of an outcome, but instead it makes an inference about the typicality. More specifically, conjunction analysis uses the proportion of the sample that shows a given outcome to infer the lower bound of typicality for that outcome (Friston et al., 1999). This inference is directly related to the statistics of the binomial distribution, which is defined as follows:

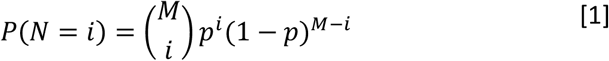

A binomial distribution with parameters *M* and *p* is the discrete probability distribution of *N* successes in a sequence of *M* independent experiments, with a success probability *p* (adapted from Wikipedia contributors (2023)).

When *p* is not known, but *M* and *N* are known, we can calculate a confidence interval for *P* (Clopper & Pearson, 1934). Fig. 2A shows the two-sided 95% confidence intervals for the success rate *P* as a function of the ratio *N*/*M* of animals showing the effect, for *M* = 1, 2, 3, 4, 5, 10 tested animals (for *M*≥10, see Fig. 4 of Clopper and Pearson (1934)). This figure nicely illustrates that increasing numbers of animals lead to decreasing widths of the confidence intervals, and an infinite number of animals would let the confidence interval shrink to the diagonal.

We now have to consider that the outcomes of statistical tests in individual subjects are imperfect and are characterized by (1) a false-positive rate, *α*, of the individual tests, and (2) a true-positive rate, *β*, of the individual tests, also referred to as sensitivity. Thus, the probability *p* of a significant statistical test is not identical to the true typicality *γ* in the population, but is a monotonic function of *γ*, parameterized by *α* and *β*:

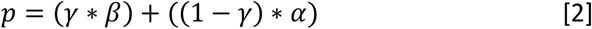

leading to

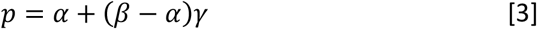

and then to

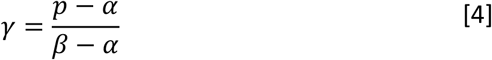

In this equation, *α* is the false-positive rate of the tests, which is typically set to 0.05. Thus, a small proportion of significant tests are false positives, slightly reducing the estimated typicality. The sensitivity *β* cannot be determined, because it is the proportion of significant tests that are truly positive, and we have no access to this truth. If we would assume that sensitivity was less than one, a nonsignificant test could reflect a false-negative test (and therefore essentially be discarded), allowing for any typicality to be consistent with the outcomes of the tests in the sample. This can be concluded directly from the formula: If we chose *β* to be less than one, the resulting lower bound *γ*_*c*_ would increase without limit, which is obviously meaningless. Therefore, we need to make the conservative assumption that sensitivity is one, as has been argued before (Friston et al., 1999). This paragraph has been adapted from Fries and Maris (2022).

By plugging the standard assumptions for *α* and *β*, and the 95% confidence intervals for the success rate *P* (see Fig. 2A), into equation [4], we obtain the two-sided 95% confidence intervals for the typicality, expressed as a function of the ratio *N*/*M* (Fig. 2B).

Because we want to be conservative, from these confidence intervals, we need to use the lower bound for typicality, which we refer to as *γ*_*c*_. Fig. 3A and Table 1 show *γ*_*c*_ as a function of *N* given different values of *M*, and reveal that *γ*_*c*_ for 2-oo-2 is merely 11%, and *γ*_*c*_ for 2-oo-3 is merely 5%. We have previously proposed that 50% is the lowest useful value for typicality, because it corresponds to the expected presence of an effect in a simple majority of the population (Fries & Maris, 2022).

**Table 1.** The lower bound (*γ*_*c*_, expressed as percentage) of the two-sided 95% confidence interval for the typicality (*γ*), as a function of the number of animals showing an effect (N) out of the number of tested animals (M), with α=0.05 and β=1.

| **M \ N** | 0 | 1 | 2 | 3 | 4 | 5 | 6 | 7 | 8 | 9 | 10 |
| --- | --- | --- | --- | --- | --- | --- | --- | --- | --- | --- | --- |
| 1 | 0 | 0 |  |  |  |  |  |  |  |  |  |
| 2 | 0 | 0 | 11.4 |  |  |  |  |  |  |  |  |
| 3 | 0 | 0 | 4.7 | 25.5 |  |  |  |  |  |  |  |
| 4 | 0 | 0 | 1.9 | 15.2 | 36.6 |  |  |  |  |  |  |
| 5 | 0 | 0 | 0.3 | 10.2 | 24.6 | 45.1 |  |  |  |  |  |
| 6 | 0 | 0 | 0 | 7.2 | 18.2 | 32.5 | 51.7 |  |  |  |  |
| 7 | 0 | 0 | 0 | 5.2 | 14.1 | 25.3 | 39.1 | 56.9 |  |  |  |
| 8 | 0 | 0 | 0 | 3.7 | 11.3 | 20.5 | 31.5 | 44.6 | 61.1 |  |  |
| 9 | 0 | 0 | 0 | 2.6 | 9.2 | 17.1 | 26.2 | 36.8 | 49.2 | 64.6 |  |
| 10 | 0 | 0 | 0 | 1.8 | 7.5 | 14.4 | 22.4 | 31.3 | 41.5 | 53.2 | 67.5 |

**Figure 3.**
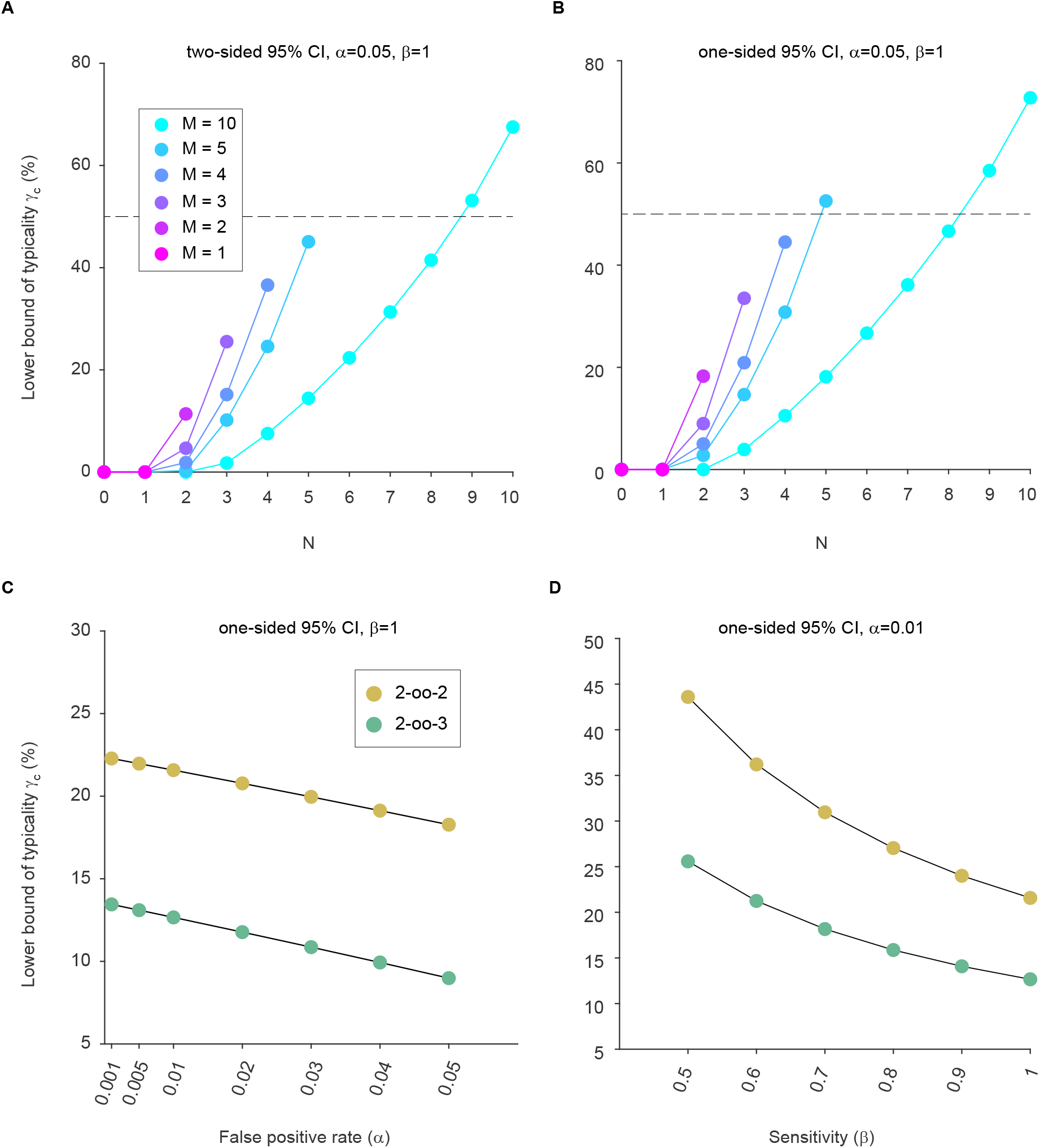
(A) The lower bound (*γ*_*c*_, expressed as percentage) of the two-sided 95% confidence interval for the typicality (*γ*), as a function of *N*, the number of animals showing an effect (i.e. an individually significant test), and *M*, the number of animals tested. The false-positive rate *α* was set to 0.05, and the sensitivity *β* was assumed to be 1, as explained in the main text. (B) Same as (A), but using one-sided 95% confidence intervals. (C) *γ*_*c*_ as a function of the false-positive rate *α*, with the sensitivity *β* fixed at a value of 1, and using one-sided 95% confidence intervals. (D) *γ*_*c*_ as a function of the sensitivity *β*, with the false-positive rate *α* fixed at a value 0.01, and using one-sided 95% confidence intervals. Higher values of *α*, like the standard value of 0.05, would lead to even lower *γ*_*c*_. This plot of *γ*_*c*_ as a function of decreasing *β* is merely to illustrate the effect, while we maintain that *β* needs to be assumed to be 1, as previously discussed (Fries & Maris, 2022; Friston et al., 1999). For (C) and (D): If we had assumed two-sided 95% confidence intervals, the values of *γ*_*c*_ would be even lower.

We so far only used two-sided confidence intervals. However, even when using one-sided confidence intervals (Fries & Maris, 2022; Friston et al., 1999), *γ*_*c*_ for 2-oo-2 is merely 18%, and *γ*_*c*_ for 2-oo-3 is merely 9%, thus far below 50% (Fig. 3B and Table 2).

**Table 2.**
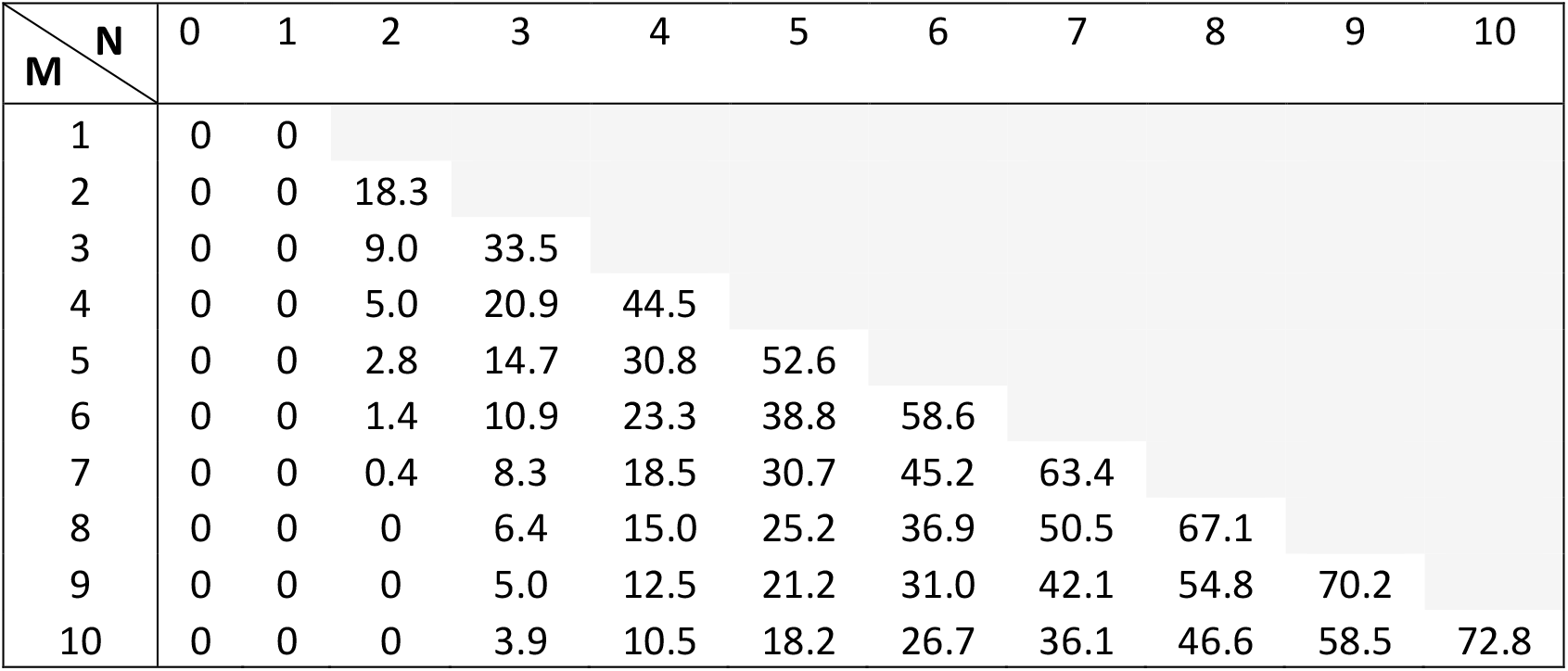
The lower bound (*γ*_*c*_, expressed as percentage) of the one-sided 95% confidence interval for the typicality (*γ*), as a function of the number of animals showing an effect (N) out of the number of tested animals (M), with α=0.05 and β=1.

Table 1 and Table 2 report the precise *γ*_*c*_ values for all possible outcomes up to N=M=10, to aid the reporting of *γ*_*c*_ values in studies that opt for this approach.

Investigators might choose a lower false-positive rate *α* for their individual tests, and this will increase the resulting *γ*_*c*_. However, even strong reductions of *α* leave *γ*_*c*_ far below 50%, for both 2-oo-2 and 2-oo-3 (Fig. 3C).

It might be argued that the assumption of *β* = 1 is necessary on theoretical grounds (see above), but that this assumption will also be wrong, because the true sensitivity will most likely never be perfect. Therefore, we explored the influence of lowering the assumption for *β*. As mentioned above, reducing *β* to arbitrarily low values will increase *γ*_*c*_ without limit and is therefore meaningless. Nevertheless, we considered reductions of *β* to a value of 0.5, which means that the test in individual subjects is so insensitive that it misses half of the subjects with an effect (or trait). Even with such strong reductions of *β*, and with *α* already reduced to 0.01, the estimate of *γ*_*c*_ remains below 50% (Fig. 3D), both for the 2-oo-2 and for the 2-oo-3 case.

Thus, the 2-oo-3 framework leads to *γ*_*c*_ values that correspond to inferences about very limited proportions of the population. Useful inferences based on 2-oo-2 or 2-oo-3 remain limited to the sample of investigated animals (Fries & Maris, 2022). Such an inference about the sample of animals is also reached with 1-oo-1. Therefore, compared to 1-oo-1, the proposal of 2-oo-2 or 2-oo-3 does not provide a gain in the quality of the inference, while at the same time doubling or even tripling the number of animals used.

### Testing null hypotheses about specific typicality values

Until now, we have only considered confidence intervals to quantify our uncertainty about typicality. Alternatively, this uncertainty can also be quantified by means of p-values. A p-value quantifies the uncertainty about a particular hypothesized typicality value, such as 50%. For a typicality of 50%, the one-sided p-value is obtained as the probability under *γ* = 0.5 of observing N or less significant outcomes. The resulting p-values are given in Table 3.

**Table 3.**
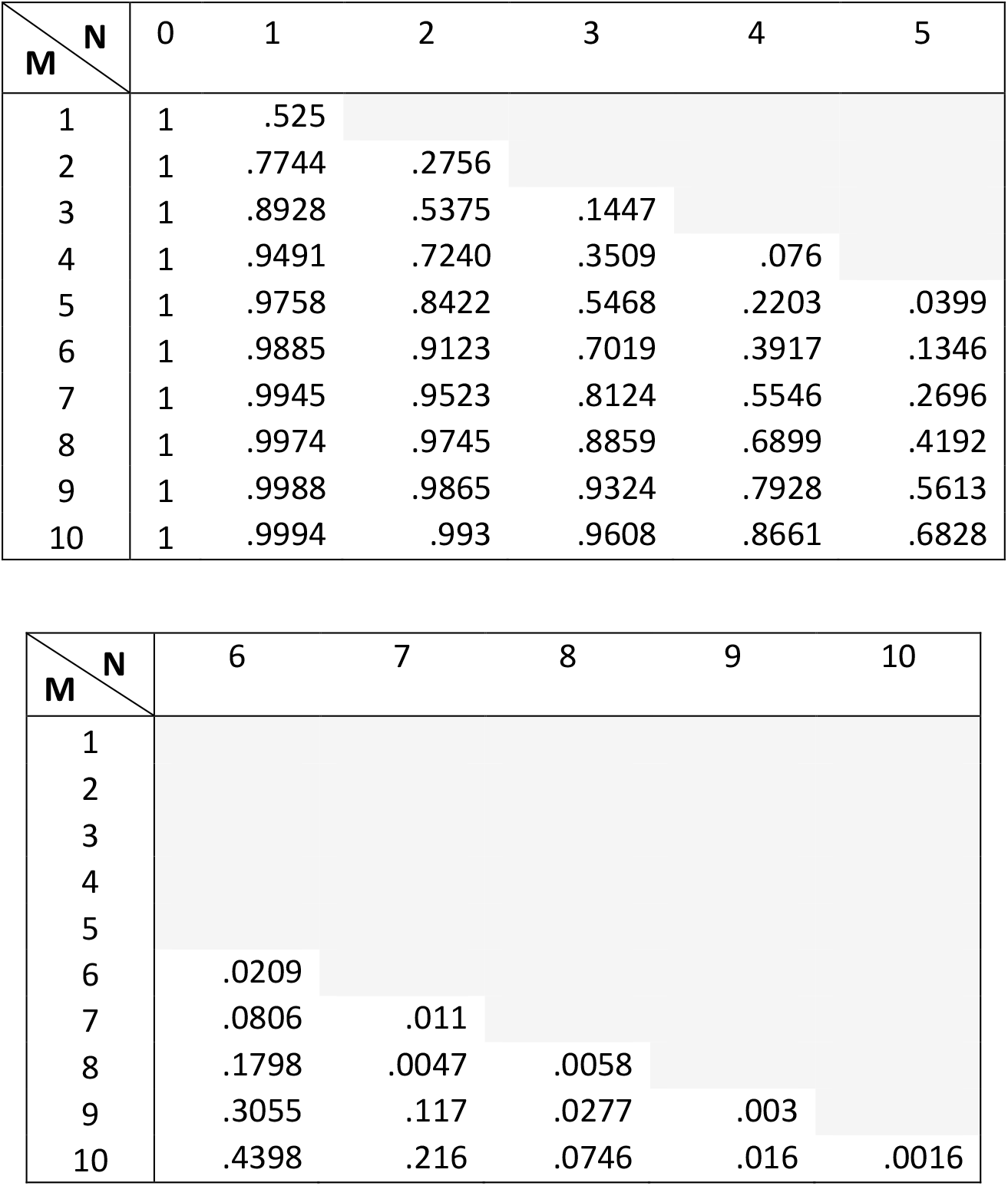
p-values for testing the majority-null hypothesis, that is, the hypothesis that the lower bound to typicality is smaller than 50%, presented as a function of the number of animals showing an effect (N) out of M tested, using α=0.05 and β=1. The top part covers cases with N ≤ 5; the bottom table covers N > 5. The p-values shown here were calculated with the code provided by (Donhauser et al., 2018), with a modification allowing the calculation for 1-oo-1.

| **M \ N** | 0 | 1 | 2 | 3 | 4 | 5 |
| --- | --- | --- | --- | --- | --- | --- |
| 1 | 1 | .525 |  |  |  |  |
| 2 | 1 | .7744 | .2756 |  |  |  |
| 3 | 1 | .8928 | .5375 | .1447 |  |  |
| 4 | 1 | .9491 | .7240 | .3509 | .076 |  |
| 5 | 1 | .9758 | .8422 | .5468 | .2203 | .0399 |
| 6 | 1 | .9885 | .9123 | .7019 | .3917 | .1346 |
| 7 | 1 | .9945 | .9523 | .8124 | .5546 | .2696 |
| 8 | 1 | .9974 | .9745 | .8859 | .6899 | .4192 |
| 9 | 1 | .9988 | .9865 | .9324 | .7928 | .5613 |
| 10 | 1 | .9994 | .993 | .9608 | .8661 | .6828 |

| **M \ N** | 6 | 7 | 8 | 9 | 10 |
| --- | --- | --- | --- | --- | --- |
| 1 |  |  |  |  |  |
| 2 |  |  |  |  |  |
| 3 |  |  |  |  |  |
| 4 |  |  |  |  |  |
| 5 |  |  |  |  |  |
| 6 | .0209 |  |  |  |  |
| 7 | .0806 | .011 |  |  |  |
| 8 | .1798 | .0047 | .0058 |  |  |
| 9 | .3055 | .117 | .0277 | .003 |  |
| 10 | .4398 | .216 | .0746 | .016 | .0016 |

### Majority-null hypothesis versus global-null hypothesis

The null hypothesis that typicality is smaller than 50% is called majority-null hypothesis. Another potential null hypothesis is that typicality is zero, i.e., that the effect does not exist in any animal. This is called global-null hypothesis. For a typicality of 0%, the one-sided p-value is obtained as the probability under *γ* = 0 of observing N or less significant outcomes. The resulting p-values are given in Table 4. Note that the p-value for 1-oo-1 is 0.05, which reflects the false-positive rate of the test conducted in that one animal. This false-positive rate can also be chosen to be lower, such that 1-oo-1 will allow to reject the global-null hypothesis with that lower false-positive rate. When we take the false-positive rate per animal into account, the same insight derives from simple logic: The null-hypothesis that the effect does not exist in any animal is rejected by observing the effect in a single animal. Note that, if the true typicality is low, testing additional animals will increase the probability for correctly rejecting the global null, i.e., it will increase the sensitivity for this rejection.

**Table 4.**
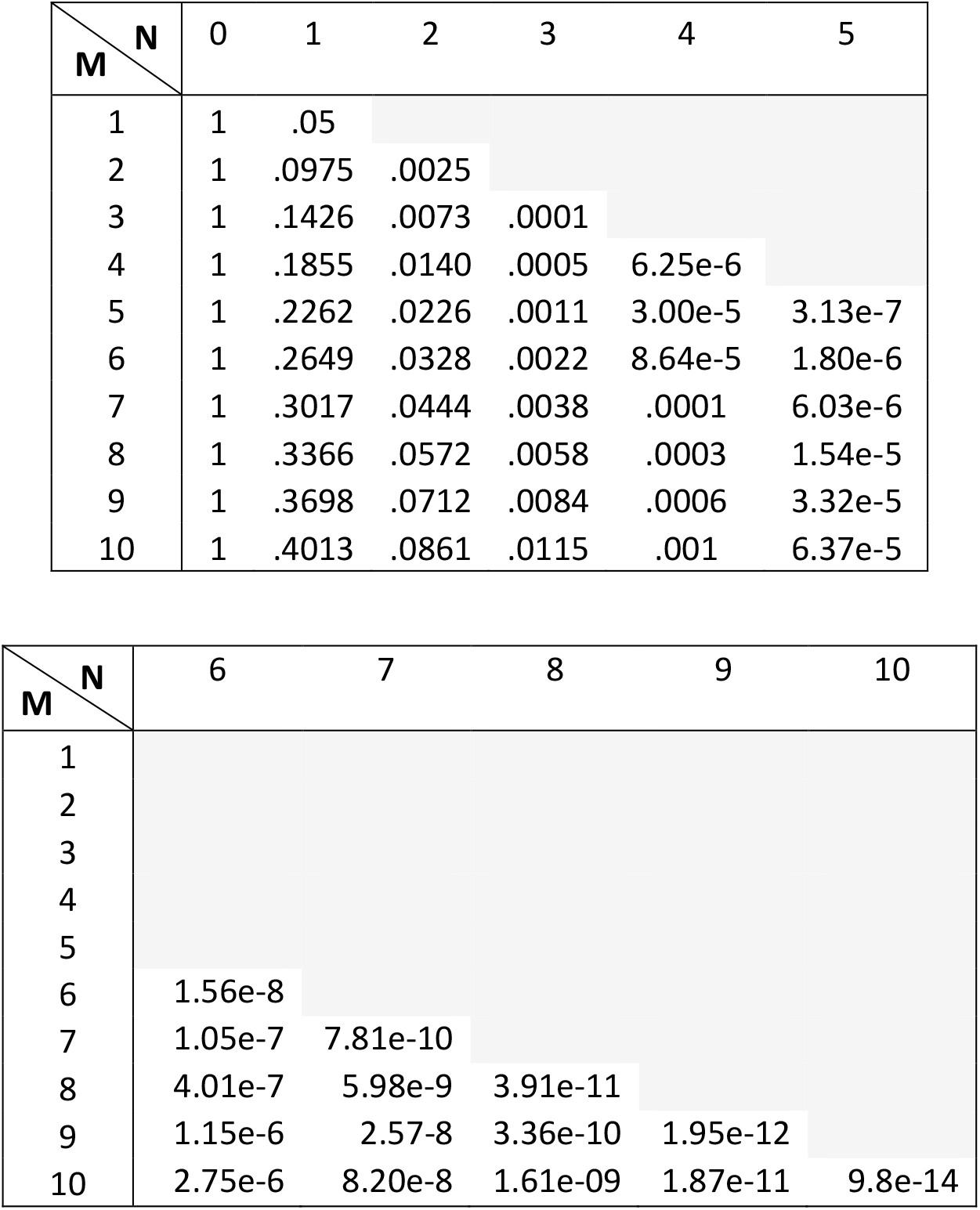
p-values for testing the global-null hypothesis, that is, the hypothesis that the lower bound to typicality is zero, presented as a function of the number of animals showing an effect (N) out of M tested, using α=0.05 and β=1. The top part covers cases with N ≤ 5; the bottom table covers N > 5. The p-values shown here were calculated with the code provided by (Donhauser et al., 2018), with a modification allowing the calculation for 1-oo-1.

Thus, the observation of an effect in few, or even a single animal, can allow the rejection of the global-null hypothesis and thereby an inference on the population. However, we maintain our previous argument that the lowest useful value of typicality for neurophysiological studies is 50%, i.e., that the useful null hypothesis is the majority-null hypothesis. The majority-null hypothesis cannot be rejected on the basis of N=1, and this typically also holds for N=2 or N=few, as we have shown here and previously (Fries & Maris, 2022). Those N-values only allow an inference on the sample. Yet, for this inference on the sample, N=1 is sufficient, and the addition of more animals is not necessary. The global-null hypothesis can be rejected on the basis of N=1, i.e., N=1 is again sufficient and the addition of one or few additional animals is not necessary.

## Conclusion

In summary, the framework proposed by Laurens (2022) has been an unconventional and welcome addition to the discussion about statistical inferences based on small numbers of animals. However, our analysis revealed that it has serious shortcomings and limitations. If studies nevertheless choose to report the individual outcomes of two or three animals, they should also report the corresponding lower bound of typicality (see Tables 1 and 2) to avoid the common misconception that the inclusion of a second or third animal would allow a general inference about the population. We maintain the previous conclusion that a useful inference about the population requires at least five animals (Fries & Maris, 2022). This number is currently not realized in typical NHP experiments. Therefore, any useful inference will remain limited to the investigated sample, and this will hold for a sample of three or two animals, or even a single animal. Consequently, we argue that the minimum required number for the publication of a typical NHP study should be one animal, to minimize the use of animals in research.

## Additional considerations

During the preparation and publication of this paper, we have received valuable comments, that we include in the following paragraphs. Some of the comments are critical arguments, which we include in quotes to make clear that it does not necessarily reflect our view (not to indicate literal quoting, as we summarize and paraphrase the arguments), and then state our respective view. Other comments are recommendations that we agree with and that we include as additional guidelines.

### Statistical standards

As mentioned in the introduction, the first and foremost point of our previous paper (Fries & Maris, 2022) and the present paper is to clarify the limitations of statistics based on two or few animals, that is, to highlight the shortcomings of traditional statistical standards in NHP (and often generally in-vivo) neurophysiology. We would greatly appreciate if those statistical standards could be improved, yet this would require the use of many more animals. Only once we have clarified this, we make the following additional point: If we accept that the number of animals cannot be increased to reach a useful population inference, then the field should embrace that the only achievable inference is on the sample, and that this inference can also be achieved with a single animal.

A prominent discussion about the absence of proper statistical standards concerns the replication crisis, caused by practices like p-hacking, researcher degrees of freedom, HARKing, selective reporting, lack of multiple comparison correction and so on (Mayo, 2018). These practices must not be applied, neither in many animals nor in a single one. The respective critique is orthogonal to and fully congruent with our highlighting of the limitations of studies with two or few animals. In fact, one of our main recommendations is that studies that do decide to report the results from two or few animals ought to report the lower bound of typicality that can actually be obtained with this approach, and thereby to transparently communicate the limitations of the respective inference. This would clarify that a 2-oo-2 or a 2-oo-3 result only allow the expectation that the next tested animal would show the effect with a probability of at least 18% or 9%, respectively. Thus, it would prevent the expectation of a guaranteed replication, and thereby help against the replication crisis. Furthermore, we have pointed out that a random-effect test with two or few animals has an extremely low sensitivity (Fries & Maris, 2022). If such low-powered random-effect tests were performed, they would be very likely to fail in an attempted replication.

### Replication as a means to make the test more severe

“A replication in a second or in few animals makes the test more severe (Mayo, 2018).” The validity of this argument depends on whether the approach is (1) a fixed-effect test on the pooled data from a sample of animals, (2) several fixed-effect tests per animal from a sample of animals, without a formal conjunction analysis, or (3) several fixed-effect tests combined in a formal conjunction analysis that provides some inference about the population. Here are the detailed considerations for each option:

(1) The severity of the test is not automatically increased by the mere addition of more animals. In NHP papers with two or few animals, the data are typically pooled over animals for a fixed-effect test. In this case, adding more animals will not change test severity. Yet, in this approach, test severity can be increased at no cost by lowering the p threshold (irrespective of using few, two or a single animal). Yet note that a higher test severity does not change the quality of the inference: The inference will remain limited to the sample.

(2) The severity of the test is actually increased if significant tests are requested per animal, which is sometimes done. However, also in this approach, a mere increase in the severity of the test does not change the quality of the inference, which still remains limited to the sample of animals.

(3) The quality of the inference could be changed if the results of multiple animals were combined in a formal conjunction analysis to draw an inference on the lower bound of typicality. Yet, the level of typicality that can be attained with two or few animals is strongly limited, as we have shown here and previously (Fries & Maris, 2022).

Importantly, the following considerations apply to approaches (2) and (3): (A) The results from all tested animals have to be included; (B) A proper control of the false-positive rate must anyhow be accomplished within each single animal, and must not require the combination of multiple independent tests. The level at which the false-positive rate is controlled per animal can be increased at the discretion of the investigators by reducing the pre-specified false-positive rate (i.e., alpha level), as mentioned already for (1); (C) The request for multiple significant tests can lead to low sensitivity (with a correspondingly high probability of failing replications). As noted above, the true value of sensitivity is unknown, must be assumed to be one for the estimation of the lower bound of typicality, yet in empirical tests is almost certainly below one. Most likely, sensitivity is higher for simple tests on fundamental effects (e.g., increase of average neuronal activity in primary sensory areas upon appropriate sensory stimulation), and lower for complicated tests on complex effects (e.g., correlation between single-trial behavioral performance and single-trial inter-areal phase alignment (Rohenkohl et al., 2018)). Crucially, if the sensitivity for each of three animals is, e.g., 0.9, then the sensitivity for the combination is 0.9^3^=0.73, i.e., the effect is actually present but missed in more than a quarter of the combined tests.

“*NHP studies often do not only report a single result, but a combination of results supporting the same conclusion. If a combination of results is replicated in a second animal, this safeguards against reporting a false positive*.”

We consider a result to be a finding that is supported by a significant statistical test. We think that a single result can be sufficient to support a conclusion and justify a publication, and this has been the case in countless publications. If the same conclusion is supported by additional results, this can provide additional evidence, but is not necessary.

Furthermore, if the replication of a full pattern is requested in a second or third animal, this further reduces sensitivity. If the sensitivity for each of three tests in a given animal is, e.g., 0.9, then the sensitivity for the combination is 0.9^3^=0.73, i.e., the effect is actually present but missed in more than a quarter of the animals. If then the replication of the full pattern is requested in a second or third animal, this reduces the sensitivity to 0.73^2^=0.53 or 0.73^3^=0.39, respectively, i.e., the effect is actually present but missed in almost half or even more than half of the combined tests.

### Consistency in experiments and data recording

Results can differ between animals due to differences between experiments or data recording. To minimize such differences, all experimental protocols including behavioral training and data recording should be as consistent as possible across the animals in a study and should be reported in great detail, including potential unavoidable differences between individuals.

### Typicality versus prevalence

In some fields of research, the term “prevalence” is used for the same quantity that we refer to as “typicality”. In those fields, “typicality” is defined as a prevalence larger than 0.5, the “majority null hypothesis” is defined as prevalence<0.5, and the “global null hypothesis” is defined as prevalence=0 (Allefeld et al., 2016; Donhauser et al., 2018; Ince et al., 2021; Rosenblatt et al., 2014); we note this here, but maintain throughout this text the meaning of “typicality” as defined above and in Friston et al. (1999) and Fries and Maris (2022).

### Fostering meta-analyses

Given that typical NHP studies report results from only two or few animals, it appears as highly useful to consider meta-analyses across individual studies. For individual studies to be useful for meta-analyses, they need to report the results of individual animals. We fully agree to this, yet we emphasize the need to distinguish between, on the one hand, the potential use of results reported in a given study for subsequent meta-analyses, and, on the other hand, the statistical inference of a given study. Regarding the statistical inference within a given study, we maintain our earlier conclusions, that two or few animals do not allow a useful inference on the population, and that existing data from two or few animals are best pooled in a fixed-effect test (Fries & Maris, 2022). This point is about a single given study, whereas the point about meta-analyses is about the potential combination of several studies. We think that the primary objective (and responsibility) of a single given study is to provide a useful inference about its respective research question; the potential additional benefit of the re-use of the results in a later meta-analysis is highly welcome, yet separate from the primary objective. For example: A study might report, as its primary result, a significant fixed-effect test on the pooled data from two animals, and as a support for future meta-analyses, the individual results of each animal. Let us assume that the individual results are significant for one animal, yet only non-significantly trending in the same direction for the second animal. This lack of significance in the second animal should not be taken as evidence against the validity of the fixed-effect test used for the primary inference of this study. Such an approach would reduce concerns about reporting the results from individual animal and thereby foster meta-analyses.

Meta-analyses would generally be fostered by the option to publish NHP datasets from single animals. We are aware of many single-monkey datasets that have so far remained unpublished, just because they have not been confirmed in a second monkey. The reasons for the lack of a second animal are often circumstantial, like the end of a work contract or of an ethics approval. If those single-monkey datasets were published, this would provide both interesting insights on the sample, and potential future insights on the population through meta-analyses.

## Author Contribution

**Eleni Psarou:** Methodology, Software, Formal analysis, Visualization, Writing - Original Draft, Writing - Review & Editing. **Christini Katsanevaki:** Methodology; Software; Formal analysis; Visualization; Writing - Original Draft; Writing - Review & Editing. **Eric Maris:** Conceptualization; Methodology; Software; Supervision; Writing - Original Draft; Writing - Review & Editing. **Pascal Fries:** Conceptualization; Methodology; Resources; Supervision; Project administration; Funding acquisition; Writing - Original Draft; Writing - Review & Editing.

## Acknowledgements

We thank Jean Laurens for helpful discussions.

